# Establishment of a reverse genetics system for SARS-CoV-2 using circular polymerase extension reaction

**DOI:** 10.1101/2020.09.23.309849

**Authors:** Shiho Torii, Chikako Ono, Rigel Suzuki, Yuhei Morioka, Itsuki Anzai, Yuzy Fauzyah, Yusuke Maeda, Wataru Kamitani, Takasuke Fukuhara, Yoshiharu Matsuura

## Abstract

Severe acute respiratory syndrome coronavirus 2 (SARS-CoV-2) has been identified as the causative agent of coronavirus disease 2019 (COVID-19). While the development of specific treatments and a vaccine is urgently needed, functional analyses of SARS-CoV-2 have been limited by the lack of convenient mutagenesis methods. In this study, we established a PCR-based, bacterium-free method to generate SARS-CoV-2 infectious clones. Recombinant SARS-CoV-2 could be rescued at high titer with high accuracy after assembling 10 SARS-CoV-2 cDNA fragments by circular polymerase extension reaction (CPER) and transfection of the resulting circular genome into susceptible cells. Notably, the construction of infectious clones for reporter viruses and mutant viruses could be completed in two simple steps: introduction of reporter genes or mutations into the desirable DNA fragments (~5,000 base pairs) by PCR and assembly of the DNA fragments by CPER. We hope that our reverse genetics system will contribute to the further understanding of SARS-CoV-2.

## Main text

Severe acute respiratory syndrome coronavirus 2 (SARS-CoV-2) in the genus *Betacoronavirus* in the family *Coronaviridae* is the causative agent of a global pandemic of severe respiratory disease, coronavirus disease 2019 (COVID-19)(Gorbalenya *et al.*, 2020). The virus was initially discovered in Wuhan, China, in late December 2019 (Zhu *et al.*, 2020; Zou *et al.*, 2020; Wu *et al.*, 2020) and has spread worldwide. As of August 26, 2020, more than 20 million COVID-19 cases have been confirmed in over 180 countries, and more than 0.8 million deaths have been reported (https://covid19.who.int/). The development of effective vaccines and specific treatments is necessary to overcome the current COVID-19 pandemic. To develop effective antivirals and safety vaccines against SARS-CoV-2, further understanding of the proliferation and pathogenesis of SARS-CoV-2 is needed. Although a large number of papers have been published since the emergence of SARS-CoV-2, neither the functions of the viral proteins nor the molecular mechanisms of propagation and pathogenesis of SARS-CoV-2 have been fully characterized. The development of a simple and efficient reverse genetics system is urgently needed for further molecular studies of SARS-CoV-2.

While a variety of infectious clones harboring the full-length viral cDNA under suitable promoters in the plasmid have been established, the plasmid system is not available for coronaviruses because of the large size of their viral genomes [−30 kilobases (kb)]. Instead, bacterial artificial chromosomes (BAC) or in vitro ligation of viral cDNA fragments have been classically utilized (Almazán *et al.*, 2006; Yount *et al.*, 2003; Scobey *et al.*, 2013; Terada *et al.*, 2019). Although these systems have allowed us to conduct molecular studies of coronaviruses, they also have some disadvantages, particularly when performing mutagenesis. In the case of BAC, undesired mutations, such as deletions or insertions, can be introduced during bacterial amplification, and verification of the full-length genome every time is time-consuming. Moreover, the in vitro ligation method is complicated and requires specific skills. Given these facts, it seems difficult to rapidly introduce reporter genes or multiple mutations into viral genes by the classical methods.

Recently, a method for the rapid generation of flavivirus infectious clones by circular polymerase extension reaction (CPER) was reported (Edmonds *et al.*, 2013). In this approach, cDNA fragments covering the full-length viral genome and a linker fragment, which encodes the promoter, polyA signal and ribozyme sequence, are amplified by PCR. Because the amplified fragments are designed to include overlapping ends with adjacent fragments, the amplified fragments can be extended as a circular viral genome with a suitable promoter by an additional PCR using the amplified fragments. By direct transfection of the circular viral genome with the promoter into susceptible cells, infectious viruses can be recovered. This means that infectious clones of flaviviruses can be constructed without any bacterial amplification or in vitro ligation. Using this CPER method, multiple reporter flaviviruses and chimeric flaviviruses have been constructed (Tamura *et al.*, 2018; Piyasena *et al.*, 2019), and a variety of mutant flaviviruses were easily generated and analyzed at the same time (Setoh *et al.*, 2019). These studies showed that CPER is an effective approach for the characterization of viral proteins.

In this study, we tried to establish a CPER method for the construction of SARS-CoV-2 recombinants possessing reporter genes and mutations. In addition, we compared the biological characteristics of the recombinants rescued by the CPER method with the parental SARS-CoV-2.

First, we examined whether the CPER approach would be applicable for the construction of an infectious clone of SARS-CoV-2. For this purpose, we used the SARS-CoV-2 strain HuDPKng19-020, which was provided by Dr. Sakuragi at the Kanagawa Prefectural Institute of Public Health. A total of 10 viral gene fragments (G1 to G10) covering the entire genome of SARS-CoV-2, and a UTR linker fragment encoding the sequences of the 3’ 43 nt of SARS-CoV-2, hepatitis delta virus ribozyme (HDVr), bovine growth hormone (BGH) polyA signal, cytomegalovirus (CMV) promoter and the 5’ 25 nt of SARS-CoV-2, were cloned into plasmids. Then, cDNA fragments of F1 to F10 and the UTR linker, possessing complementary ends with 25 to 452 overlapping nucleotides, were amplified with specific primers and subjected to CPER as templates (Figure 1A and Table S1). The cDNA fragments of F9 and F10 were connected to F9/10 before CPER by overlap PCR. A negative control was prepared by CPER using cDNA fragments excluding F9/10. The full-length cDNA clone of SARS-CoV-2 under the CMV promoter obtained by CPER (condition 1 described in Star methods) was directly transfected into either BHK-21 cells or tetracycline-inducible ACE2 and TMPRSS-expressing IFNAR1-deficient HEK293 (HEK293-3P6C33) cells without any purification steps. Because the SARS-CoV-2 nucleocapsid protein was reported to enhance the propagation of coronavirus RNA transcripts (Xie *et al.*, 2020), the nucleocapsid-expressing plasmid was transfected together with the CPER products into cells. Upon the induction of ACE2 and TMPRSS expression in HEK293-3P6C33 cells or the overlaying of Vero cells expressing TMPRSS2 (VeroE6/TMPRSS2) onto BHK-21 cells, propagation of SARS-CoV-2 was assessed by cytopathic effects (CPE).

**Figure 1.**
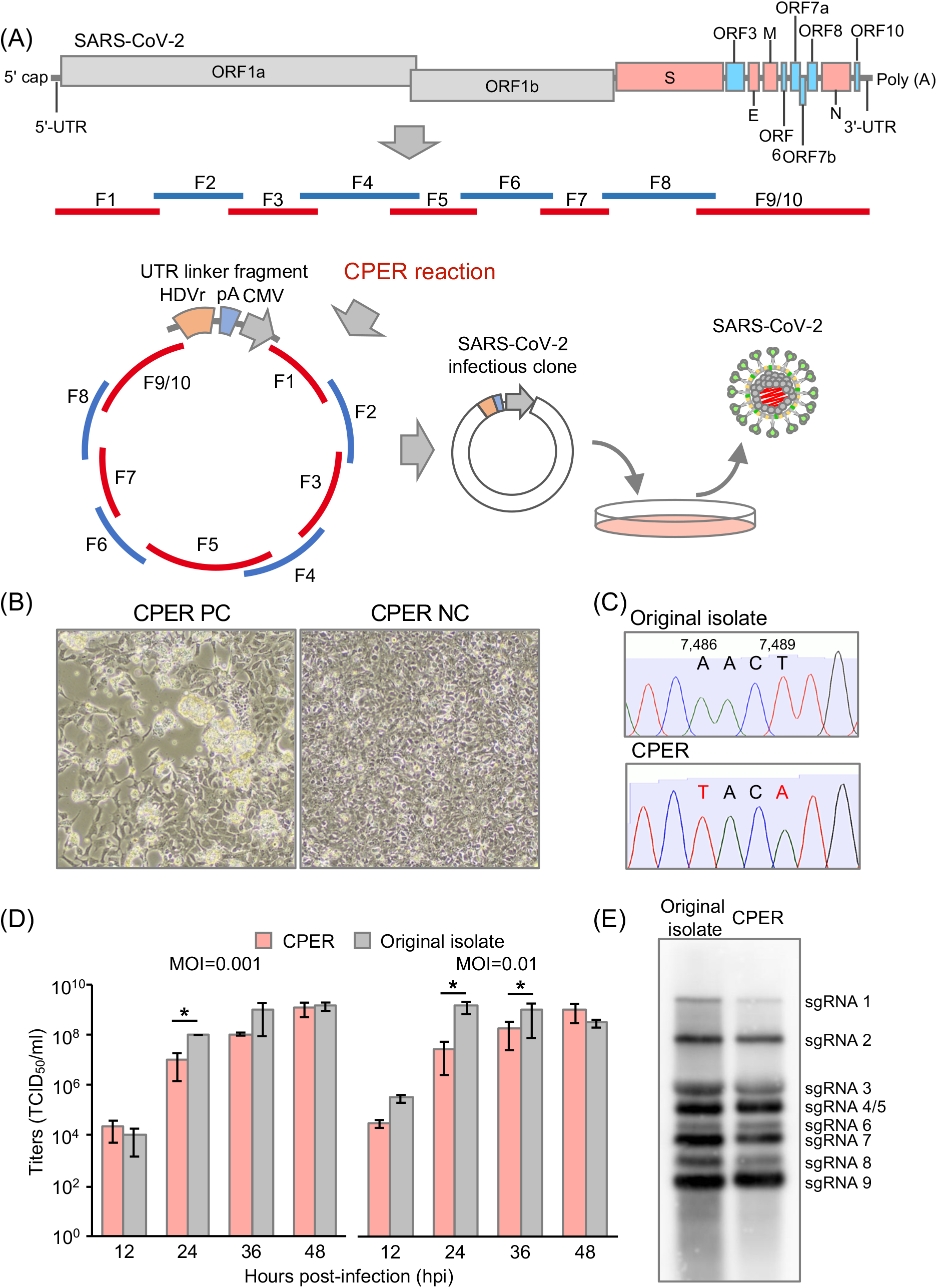
Establishment of CPER-based reverse genetics for SARS-CoV-2. (A) Schematic representation of a CPER approach for the generation of recombinant SARS-CoV-2. A total of 9 fragments (F1to F8, and F9/10) covering the full-length of the SARS-CoV-2 genome were amplified, then assembled with a UTR linker fragment including the HDVr, the BGH polyA signal and the CMV promoter by CPER. The resulting CPER products were transfected into the susceptible cells. (B) HEK293-3P6C33 cells were transfected with the CPER product and the bright field image was acquired at 7 days post-transfection (dpt) (left). As a negative control, the CPER product obtained without fragment F9/10 was transfected into cells and the bright field image was obtained at 7 dpt (right). (C) Genetic markers (2 silent mutations, A7,486T and T7,489A) in the recombinant SARS-CoV-2 genome. (D) Comparison of the growth kinetics of the recombinant SARS-CoV-2 generated by CPER with that of the original isolate. VeroE6/TMPRSS2 cells were infected with the viruses (MOI=0.001 or 0.01), and infectious titers in the culture supernatants of the SARS-CoV-2-infected cells were determined by TCID_50_ assay from 12 to 48 hours post-infection (hpi). (E) Northern blot analyses of subgenomic RNAs. RNAs extracted from cells infected with the parental virus and the recombinant SARS-CoV-2 recovered by CPER were subjected to Northern blot analyses.

At 7 days post-transfection (dpt), CPE were observed in HEK293-3P6C33 cells transfected with the CPER product (CPER PC), but not in those transfected with the negative control (CPER NC) (Figure 1B). CPE was not observed in BHK-21cells until 14 dpt of the CPER products (data not shown). Propagation of the progeny viruses was examined by serial passages of the culture supernatants of HEK293-3P6C33 cells in VeroE6/TMPRSS2 cells for 2 rounds. CPE were observed at 1 day post-infection (dpi) of the viruses of both Passage 1 (P1) and P2 (data not shown). Infectious titers of the culture supernatants of HEK293-3P6C33 cells at 7 dpt (P0), P1 and P2 viruses, collected at 2 dpi to VeroE6/TMPRSS2 cells, were determined by 50% tissue culture infective dose (TCID_50_) assays. The infectious titers of the P0, P1, and P2 viruses were 10^5.8^, 10^6.3^, and 10^5.8^ TCID_50_/ml, respectively (data not shown), demonstrating that infectious SARS-Co2V-2 was rescued at high titer upon transfection of the CPER product into HEK293-3P6C33 cells, and the recovered viruses were capable of propagating well in VeroE6/TMPRSS2 cells.

To optimize the conditions of recovery of infectious SARS-CoV-2 particles by CPER, the reactions were performed using different numbers of cycles, steps and extension times (conditions 1 to 3 in Star Methods). To investigate the effect of expression of nucleocapsid as shown previously (Xie *et al.*, 2020) on the recovery of infectious particles, CPER products were transfected into HEK293-3P6C33 cells with or without an expression plasmid of the nucleocapsid protein. Culture supernatants of HEK293-3P6C33 cells transfected with the CPER product were collected at the indicated time points for 9 days, and infectious titers were determined as the TCID_50_. In cells transfected with the CPER products without F9/10 (CPER NC), no CPE and no infectious titer in the supernatants was detected until 9 dpt (condition 3 without the expression plasmid of the nucleocapsid protein is shown in Figure S1A). On the other hand, infectious titers were detected from 5 dpt and reached around 10^7.0^ TCID_50_/ml in the supernatants of cells transfected with the CPER products, regardless of the reaction conditions (condition 3 without the expression plasmid of the nucleocapsid protein is shown in Figure S1A). No effect of the expression of nucleocapsid protein was observed, suggesting that nucleocapsid is not necessary to recover infectious particles in this method. We selected condition 3 (an initial 2 minutes of denaturation at 98°C; followed by 35 cycles of 10 seconds at 98°C, 15 seconds at 55°C, and 15 minutes at 68°C; and a final extension for 15 minutes at 68°C) for further CPER to generate an infectious cDNA clone for the recovery of infectious particles after transfection into HEK293-3P6C33 cells. Condition 3 was chosen because CPE appeared in cells at 5 dpt of CPER products obtained by condition 3, but at 7 dpt of those obtained by conditions 1 and 2 (data not shown).

To determine the full-length genome sequences of viruses recovered by the CPER method, 2 viruses (#1 and #2 in Table S2), which were obtained independently at different time points from the supernatants of HEK293-3P6C33 cells, were passaged two times in VeroE6/TMPRSS2 cells, and subjected to Sanger sequence analysis with specific primers. Sequence analyses of P0–P2 viruses demonstrated that the recombinant viruses maintained genetic markers (two silent mutations, A7486T and T7489A; Figure 1C), indicating that there was no contamination of parental virus. Importantly, except for the genetic markers, there was only one difference (T to T/A) in all tested P0 viruses, suggesting that the reverse genetic system for SARS-CoV-2 by CPER had high accuracy. While a large deletion occurred in P1 and P2 of the #2 virus, that deletion was reported to occur during passage in Vero E6 cells (Lau *et al.*, 2020).

Next, we investigated the growth kinetics of recombinant viruses in comparison with parental SARS-CoV-2. Recombinant viruses, which were recovered at 7 dpt of CPER products into HEK293-3P6C33 cells, and parental SARS-CoV-2 were infected into VeroE6/TMPRSS2 cells at multiplicities of infection (MOIs) of 0.001 and 0.01. Infectious titers in the culture supernatants of VeroE6/TMPRSS2 cells were determined for 48 hours. Although the rescued viruses by CPER (CPER) exhibited lower titers than the parental viruses (Original isolate) at 24 and 36 hours post-infection (hpi), no significant difference in maximum titers was observed between the rescued and parental viruses (Figure 1D). These results suggest that propagation of the rescued SARS-CoV-2 by CPER is slow but reached levels similar to those reached by the parental virus, as previously reported (Xie *et al.*, 2020). To examine the viral RNA synthesis of the rescued viruses, Northern blot analyses were performed. In total, eight subgenomic RNAs were detected in cells infected with both parental and rescued viruses, and all eight RNAs were similar in size (Figure 1E). Taken together, these results showed that SARS-CoV-2 rescued by the CPER method exhibits biological characteristics similar to those of the parental virus.

Next, we applied the CPER method for construction of recombinant SARS-CoV-2 carrying reporter genes. The nucleotide sequences from 27,433 to 27,675 in ORF7 were replaced by the sfGFP gene, as previously reported (Thi Nhu Thao *et al.*, 2020) (Figure 2A). Using the DNA fragments containing the sfGFP gene, infectious DNA clones were assembled by CPER. CPE was observed in HEK293-3P6C33 cells at 7 dpt of CPER product, and the insertion of the reporter genes in the viruses recovered in the culture supernatants was confirmed by Sanger sequence analysis (data not shown). Then, the wild-type (WT) virus and sfGFP-carrying virus (GFP virus) were infected to VeroE6/TMPRSS2 cells at an MOI of 0.001 and the growth kinetics of the viruses were evaluated. GFP viruses exhibited lower titers than WT viruses at the indicated time points and the maximum titers of GFP viruses at 36 hpi were significantly lower than those of WT viruses at 48 hpi (Figure 2B). We also examined the expression of GFP in Vero-TMPRSS2 cells upon infection with the GFP recombinant from 12 to 36 hpi by fluorescence microscopy. The numbers of GFP-positive cells were increased from 12 to 36 hpi and almost all cells exhibited the fluorescent signals at 24 and 36 hpi (Figure 2C). These results indicate that recombinant SARS-CoV-2 harboring reporter genes could be quickly engineered by the CPER method and the GFP viruses demonstrated lower growth kinetics than the WT viruses.

**Figure 2.**
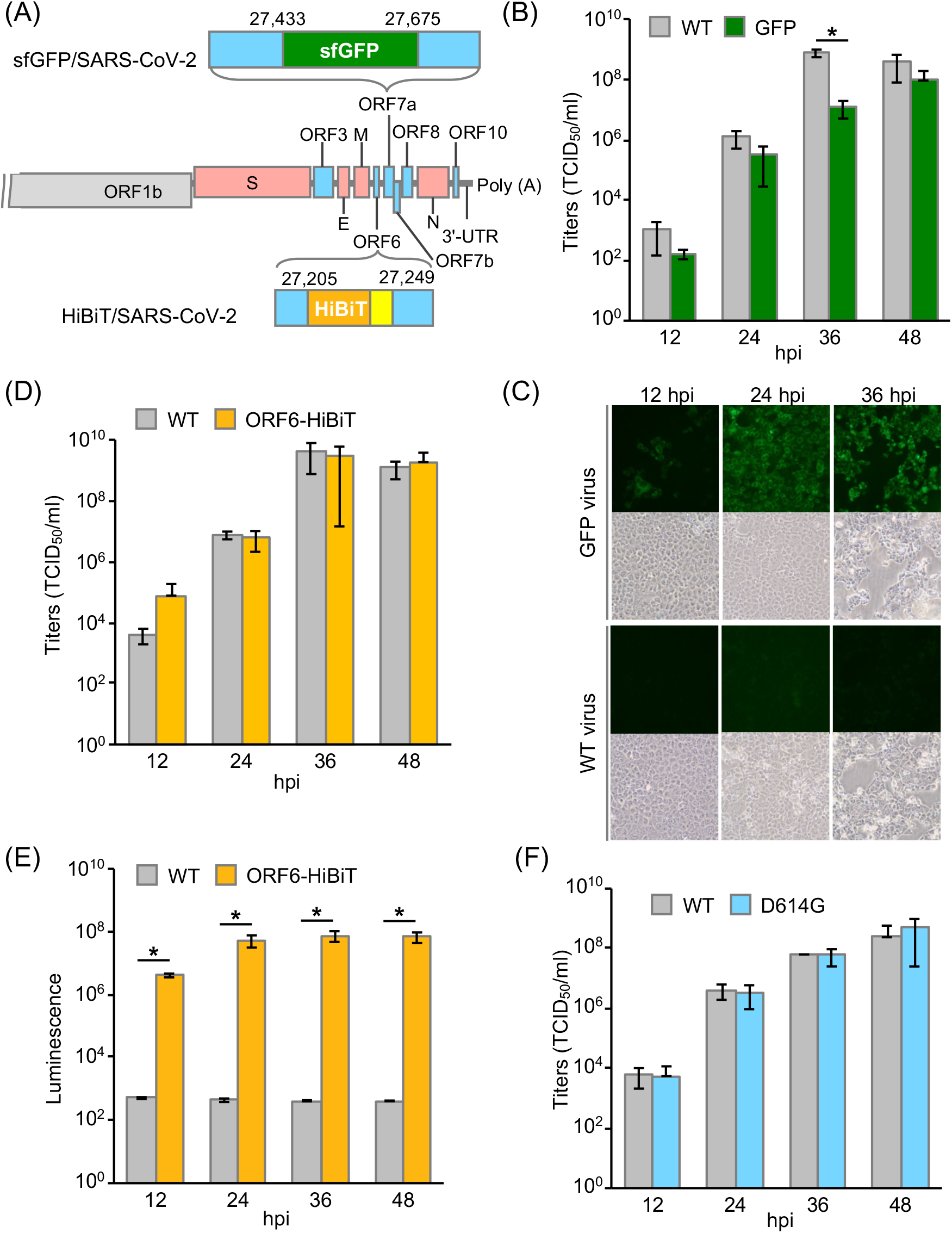
Characterization of SARS-CoV-2 recombinants possessing reporter genes and mutations. (A) Gene structure of recombinant SARS-CoV-2 carrying the sfGFP or HiBiT gene. The nucleotide sequences of 27,433–27,675 in ORF7 were replaced with those of sfGFP. The HiBiT gene was inserted into the N terminus of ORF6. (B) Growth kinetics of wild-type SARS-CoV-2 (WT) and SARS-CoV-2 carrying sfGFP (GFP). VeroE6/TMPRSS2 cells were infected with the viruses (MOI=0.001), and infectious titers in the culture supernatants were determined at the indicated time points. (C) The fluorescent signal in VeroE6/TMPRSS2 cells infected with the WT virus and GFP virus was observed for 36 hpi. (D) Growth kinetics of the WT virus and recombinant virus possessing the HiBiT gene in ORF6 (ORF6-HiBiT). Titers in the culture supernatants of VeroE6/TMPRSS2 cells, infected with the viruses (MOI=0.01), were measured for 48 hours. (E) Luciferase activities in VeroE6/TMPRSS2 cells infected with the WT virus and ORF6-HiBiT virus were determined from 12 to 48 hpi. (F) Infectious titers in the culture supernatants of VeroE6/TMPRSS2 cells infected with the WT virus or mutant virus, harboring a substitution of D614 to G in the spike protein (D614G), were determined at the indicated time points.

We also employed NanoLuc binary technology (NanoBiT) (Dixon *et al.*, 2016) in this study. NanoBiT is a split reporter consisting of 2 subunits, high-affinity NanoBiT (HiBiT) (Schwinn *et al.*, 2018) and large NanoBiT (LgBiT). By interaction of HiBiT and LgBiT in cells or in vitro, NanoLuc enzymatic activity can be detected. Because a reporter virus can be generated by inserting a small cDNA fragment encoding 11 amino acids into the viral genome, the effect on viral growth and pathogenicity has been shown to be minimum (Tamura *et al.*, 2018). In this study, we inserted a HiBiT luciferase gene (VSGWRLFKKIS) and a linker sequence (GSSG) into the N terminus of the ORF6 gene of SARS-CoV-2 (Figure 2A) by overlap PCR using DNA fragment F9 and specific primer sets (see Star Methods for details), and the recombinant SARS-CoV-2 possessing HiBiT was generated by CPER using the resulting fragments. We confirmed the incorporation of the HiBiT gene into the recombinant viruses by Sanger sequence analysis (Figure S1B). Infectious titers in the culture supernatants of VeroE6/TMPRSS2 cells infected with the HiBiT recombinants (ORF6-HiBiT) were comparable to those in the culture supernatants of the same cells infected with the WT (Figure 2D). Luciferase activities in cells infected with the recombinant, as assessed by addition of LgBiT protein into the cell lysates, were increased from 12 to 48 hpi (Figure 2E). These results suggest that recombinant SARS-CoV-2 carrying HiBiT could be easily generated by overlap PCR for introduction of HiBiT into the indicated places, followed by CPER to assemble the resulting DNA fragments, which function as reporter viruses exhibiting growth kinetics comparable to the WT virus.

Finally, we examined whether the CPER method can be applied for the construction of recombinant SARS-CoV-2 with the desired mutations. Substitution of D614 to G on the spike protein was frequently detected, and became dominant in European, South and North American countries (Koyama *et al.*, 2020) (https://www.gisaid.org/epiflu-applications/hcov-19-genomic-epidemiology/). To evaluate the effect of this substitution on virus propagation, we introduced this mutation in fragment F8 of SARS-CoV-2 (strain HuDPKng19-020), and generated the infectious clone by the CPER method. We confirmed the substitution in the recombinant by Sanger sequence analysis (data not shown), but there was no difference in growth kinetics between the WT and D614G recombinant in VeroE6/TMPRSS2 cells (Figure 2F). Collectively, these results suggest that our novel reverse genetic system based on CPER is an efficient method for quick generation of recombinants for SARS-CoV-2.

Reverse genetics is one of the essential tools to analyze the functions of viral proteins; however, due to the large size of the coronavirus genome, gene manipulations for coronaviruses have been performed by only a very limited number of groups. Here, we established a simple and quick reverse genetics system for SARS-CoV-2 based on the CPER method. The system consists of two steps—namely, amplification of fragments encoding promoter and viral genes, followed by assembly into infectious cDNA clones by PCR without any bacterial amplification. Recombinant viruses were generated with high titers at 7 dpt of CPER products into cells. Using high-fidelity polymerase, SARS-CoV-2 could be rescued with high accuracy, and the rescued virus exhibited characteristics similar to those of the parental virus. It is worth noting that the same tools, i.e., primers and promoter fragments, can be used for generation of viruses possessing multiple mutations or reporters. This method will allow us to conduct high-throughput mutagenesis of SARS-CoV-2, to clarify the function of viral proteins and the mechanisms of propagation and pathogenesis.

In this study, we first report that the CPER method is available to assemble a large size genome (30 kb) of SARS-CoV-2 with high accuracy. While infectious clones were recovered by all CPER conditions we examined, recombinant viruses were recovered only in HEK293-3P6C33 cells (Figure 1B), indicating that the transfection efficiency of the CPER product and replication efficiency of viruses might be important to produce recombinant viruses. Multiple primer sets for the generation of DNA fragments (F1–F10) were examined (data not shown); however, recombinant viruses were recovered only when using the fragments generated by the indicated primer sets (Table S1). As each SARS-CoV-2 subgenomic RNA contains a common leader sequence (72 nucleotides) at the 5’ end (Kim *et al.*, 2020), the design of optimal primer sets seemed to be important for efficient assembly of the circular genome. In the case of the CPER method for flaviviruses, it was reported that insect-specific flaviviruses, which can replicate only in insect-derived cells, could be rescued using mosquito-derived C6/36 cells by replacing the CMV promoter with the OpIE2 promoter gene in the UTR linker fragment (Piyasena *et al.*, 2017). Deletion of 23 nucleotides in the 3’ region of the promoter, enhanced the efficiency of recovery of insect-specific flaviviruses (Piyasena *et al.*, 2017). In that sense, further investigation of the CPER conditions (i.e., promoter sequences and primer sets) for SARS-CoV-2 may enhance the recovery of recombinants, even for slowly growing viruses. Moreover, it might be possible to generate other coronaviruses and chimeric coronaviruses by using susceptible cell lines and optimal promoters.

While the GFP recombinant virus exhibited a lower titer than the WT viruses, the HiBiT recombinants exhibited growth kinetics similar to that of the WT virus, suggesting that the HiBiT virus might be suitable for functional analyses. As we previously showed, HiBiT recombinant viruses could be useful for drug screening and animal experiments (Tamura *et al.*, 2019; Tamura *et al.*, 2018). By combining the reporter assay and mutational scanning, we were able to investigate the biological features of viral proteins of SARS-CoV-2.

To generate avirulent strain for use in developing a safe live-attenuated vaccine for SARS-CoV-2, the CPER method is very useful. To characterize escape mutants occurring through the use of antiviral drugs or vaccinations, CPER can be a robust tool. In comparison with previous reverse genetics systems for coronaviruses, the CPER method is easier and quicker, especially when applied for mutagenesis. We believe that this method will be widely utilized in order to investigate SARS-CoV-2 and further clarify its pathogenesis and propagation.

## Acknowledgements

We thank M. Tomiyama for her secretarial work and M. Ishibashi, and K, Toyoda, for their technical assistance. We also thank J. Sakuragi in Kanagawa Prefectural Institute of Public Health for providing SARS-CoV-2. This work was supported from the Ministry of Health, Labor and Welfare of Japan and the Japan Agency for Medical Research and Development (AMED; http://www.amed.go.jp/) (JP20wm0225002, JP20he0822006, JP20fk0108264, JP20he0822008, JP20wm0225003, JP20fk0108267, JP19fk0108113, and JP20wm0125010), and the Japan Society for the Promotion of Science KAKENHI (JP19K24679). S. Torii is supported by a JSPS Research Fellowships for young scientists (https://www.jsps.go.jp/english/e-grants/) (19J12641).

**We have no conflicts of interest to declare.**

## Author contributions

S.T. performed all experiments and analyzed the data under the supervision of T.F. and Y.M. (Matsuura); C.O., R.S., Y.M. (Morioka), I.A., Y.F., W.K., and Y.M. (Maeda) provided critical resources and scientific insights; S.T. and T.F. wrote the original draft; C.O., R.S., Y.M. (Morioka), I.A., Y.F., W.K., Y.M. (Maeda), and Y.M. (Matsuura) wrote, reviewed, and edited the final manuscript.

## Star methods

### Lead Contact

Further information and requests for resources and reagents should be directed to and will be fulfilled by either of the two Lead Contacts, Takasuke Fukuhara or Yoshiharu Matsuura.

## Materials Availability

All unique materials generated in this study are available from either of the two lead contacts with a completed Materials Transfer Agreement.

## Data Code and Availability

This study did not generate any unique datasets or code.

## Experimental Model and Subject details

### Viruses and Cells

SARS-CoV-2 strain HuDPKng19-020 was kindly provided by Dr. Sakuragi at the Kanagawa Prefectural Institute of Public Health. All viruses were initially amplified in Vero E6 cells and the culture supernatants were harvested and stored at −80°C until use. BHK-21 cells and Vero E6 cells were maintained in high-glucose Dulbecco’s Modified Eagle Medium (DMEM) (Nacalai Tesque) containing 10% fetal bovine serum (FBS) (Sigma), 100 units/ml penicillin and 100 μg/ml streptomycin (P/S) (Sigma). TMPRSS2-expressing Vero E6 (VeroE6/TMPRSS2) cells were obtained from the Japanese Collection of Research Bioresources Cell Bank (JCRB1819), and maintained in DMEM containing 10% FBS and G418 (Nacalai Tesque). IFNAR1-deficient HEK293 cells, in which human ACE2 and TMPRSS2 are induced by tetracycline, were established and designated as HEK293-3P6C33 cells. In brief, ACE2 carboxyl terminally tagged with BFP followed by IRES-TMPRSS2 was constructed under a tetracycline-responsible promoter in a piggyback-based vector (Yusa *et al.*, 2009) (kindly provided from Welcome Trust Sanger Institute). EF-1alpha-driven rtTA-P2A-bsd was also inserted into this vector. Then, this plasmid was co-transfected with a transposase-expression vector, pCMV-hyPBase (Yusa *et al.*, 2011) (kindly provided from Welcome Trust Sanger Institute), into HEK293 cells, whose IFNAR1 had been knocked out using a CRISPR/Cas9 system. The HEK293-3P6C33 cells were maintained in DMEM containing 10% FBS and Blasticidin (solution) (10 μg/ml) (Invivogen), and the exogenous expression of ACE2 and TMPRSS2 was induced by addition of doxycycline hydrochloride (1 μg/ml) (Sigma). All the above cells were cultured at 37°C under 5% CO2. All experiments involving SARS-CoV-2 were performed in biosafety level-3 laboratories, following the standard biosafety protocols approved by the Research institute for Microbial Diseases at Osaka University.

## Method details

### RNA extraction, cDNA synthesis, and determination of nucleotide sequences of the full-length SARS-CoV-2 genome

Total RNA was extracted from the supernatants of SARS-CoV-2-infected cells by using a PureLink RNA Mini Kit (Invitrogen). Thereafter, first-strand cDNA was synthesized by using a PrimeScript RT reagent kit (Perfect Real Time) (TaKaRa Bio) with random hexamer primers and extracted RNA, according to the manufacturer’s protocols. To determine the nucleotide sequences of full-length SARS-CoV-2, a total of 10 gene fragments (G1–G10), which are up to 4,000 base pairs in length and cover the entire SARS-CoV-2 sequence, were amplified with synthesized cDNA, specific primer sets for SARS-CoV-2 and PrimeSTAR GXL DNA polymerase (TaKaRa Bio). The 5’ termini of RNA were amplified by using the 5’RACE System for Rapid Amplification of cDNA Ends, Version 2.0 (Thermo Fisher Scientific) with specific primers (CoV-2-Race1 and CoV-2-Race2) as previously described (Li *et al.*, 2005). Then, the amplified products were directly sequenced in both directions by using the ABI PRISM 3130 Genetic Analyzer (Applied Biosystems) with specific primers.

### Plasmids

A total of 10 SARS-CoV-2 (HuDPKng19-020) cDNA fragments (G1–G10) were amplified by PrimeSTAR GXL DNA polymerase and cloned into plasmids. SARS-CoV-2 fragments (G1, G2, G3, G4, G7, G8, G9 and G10) were cloned into the lentiviral vector pCSII-EF-RfA (pCSII-CoV-2-G1, -G2, -G3, -G4, -G7, -G8, -G9 and -G10), and other fragments (G5 and G6) were cloned into the pMW119 vector (pMW-CoV-2-G5 and -G6). A UTR linker for SARS-CoV-2 was also generated using the pMW119 vector, which encodes sequences of the 3’ 43 nt of SARS-CoV-2, BGH polyA signal, HDVr, CMV promoter and the 5’ 25 nt of SARS-CoV-2 (pMW-CoV-2-UTRlinker). To distinguish between the recombinant viruses and the original virus, two silent mutations, A7,486T and T7,489A, were introduced into pCSII-CoV-2-G4 as genetic markers. To generate reporter viruses possessing the sfGFP gene, the nucleotide sequences of 27,433–27,675 in the ORF7 region in pCSII-CoV-2-G10 were replaced with the sfGFP gene as previously described (Thi Nhu Thao *et al.*, 2020). The resulting vectors were designated as pCSII-CoV-2-G10-sfGFP. A plasmid expressing the SARS-CoV-2 nucleocapsid was constructed by inserting the cDNA of the nucleocapsid into pCAGGS (pCAGGS-CoV-2-N). Sequences of all the inserted DNAs were confirmed by sequencing (ABI PRISM 3130 Genetic Analyzer).

### CPER reaction

SARS-CoV-2 recombinants were generated by CPER as described previously (Setoh *et al.*, 2017), with some modifications. To amplify all the cDNA fragments having complementary ends with a 25- to 452-nucleotide overlap for CPER, plasmids encoding SARS-CoV-2 gene fragments (G1–G10) and UTRlinker were used as templates. The specific primers used to amplify DNA fragments (F1-F10 and the UTR linker) are described in the Key resources table. Then, the DNA fragments of F9 and F10 were connected before CPER by overlap PCR with a primer set (CoV-2-F9-Fw and CoV-2-F10-Rv). By using equimolar amounts (0.1 pmol each) of the resulting 10 DNA fragments (F1 to F8, F9/10 and the UTR linker) and 2 μl of PrimeStar GXL DNA polymerase, CPER was performed within 50 μl reaction volumes. The cycling conditions of CPER were as follows: condition 1 (an initial 2 minutes of denaturation at 98°C; 20 cycles of 10 seconds at 98°C, 15 seconds at 55°C, and 25 minutes at 68°C; and a final extension for 25 minutes at 68°C), condition 2 (an initial 2 minutes of denaturation at 98°C; 35 cycles of 10 seconds at 98°C and 15 minutes at 68°C; and a final extension for 15 minutes at 68°C), or condition 3 (an initial 2 minutes of denaturation at 98°C; 35 cycles of 10 seconds at 98°C, 15 seconds at 55°C, and 15 minutes at 68°C; and a final extension for 15 minutes at 68°C).

The infectious clones of SARS-CoV-2 carrying sfGFP were also assembled by CPER using pCSII-CoV-2-G10-sfGFP as templates to acquire DNA fragment F10. The HiBiT recombinant SARS-CoV-2 was also generated by CPER as previously described (Tamura *et al.*, 2018), with some modifications. The HiBiT gene (VSGWRLFKKIS) and a linker sequence (GSSG) were inserted in the N terminus of the ORF6 sequence by overlap PCR using DNA fragment F9 and a specific primer set (ORF6-HiBiT-Rv and ORF6-HiBiT-Fw). Thereafter, CPER was conducted using the resulting fragment F9 containing the HiBiT gene.

### Transfection

The CPER products (25 μl out of a 50 μl reaction volume) were transfected into HEK293-3P6C33 cells or BHK-21 cells with Trans IT LT-1 (Mirus), following the manufacturer’s protocols. At 6 hours post-transfection, the culture supernatants of HEK293-3P6C33 cells were replaced with DMEM containing 2% FBS and doxycycline hydrochloride (1 μg/ml), and BHK-21 cells were overlaid by VeroE6/TMPRSS2 cells.

### Titration and growth kinetics

The infectious titers in the culture supernatants were determined by the 50% tissue culture infective doses (TCID_50_). The culture supernatants of cells were inoculated onto VeroE6/TMPRSS2 cells in 96-well plates after ten-fold serial dilution with DMEM containing 2% FBS, and the infectious titers were determined at 72 hours post-infection (hpi). For growth kinetics, SARS-CoV-2 was inoculated into VeroE6/TMPRSS2 cells in 6-well plates at a multiplicity of infection (MOI) of 0.001 or 0.01 and the culture supernatants were replaced with new media at 1 hpi and incubated for 48 hours. The infectious titers in the culture supernatants of cells collected at 12, 24, 36 and 48 hpi were determined.

### Northern blotting

Total RNAs were extracted from cells infected with the WT or recombinant SARS-CoV-2 and subjected to northern blot analysis as previously described (Xie *et al.*, 2020). A digoxigenin (DIG)-labeled random-primed probe, corresponding to 28,999 to 29,573 of the SARS-CoV-2 genome, was generated by using a DIG RNA Labeling kit (SP6/T7) (Roche), and utilized to detect viral mRNAs. The RNAs were washed with the DIG luminescent detection kit (Roche) and visualized with CDP-Star Chemiluminescent Substrate (Roche), according to the manufacturer’s protocols.

### HiBiT luciferase assay

SARS-CoV-2 infected cells were collected at the indicated time points and subjected to luciferase assay. Luciferase activity was measured by using a Nano-Glo HiBiT Lytic assay system (Promega), following the manufacturer’s protocols. In brief, the HiBiT assay was conducted by adding Nano-Glo substrate and LgBiT protein into the lysates of cells infected with viruses, and then measuring the luciferase activities with a luminometer.

### Statistical analysis and normalizing

All assays were performed independently at least 2 times. The data were expressed as means ±S.D. Statistical significance was determined by the two-tailed Student’s *t*-test. *P*-values <0.05 were considered significant and indicated by a single asterisk (*).

## The supplemental information includes 1 figure and 2 tables

**Figure S1. Characterization of recombinant SARS-CoV-2.**

(A) Time course analysis of infectious SARS-CoV-2 production. CPER products were transfected into HEK293-3P6C33 cells (CPER PC), and infectious titers in the culture supernatants were measured at the indicated time points. As a negative control, the CPER product obtained without fragment F9/10 was transfected into cells (CPER NC). (B) Sequence analysis of the recombinant virus possessing the HiBiT gene in ORF6. The HiBiT gene and a linker sequence were inserted into the N terminus of the ORF6 sequence.

**Table S1. SARS-CoV-2 DNA fragments used for CPER reaction**

**Table S2. Mutations of recombinant SARS-CoV-2 (P0–P2 viruses)**

