## Supplemental figures for "Establishment of a reverse genetics system for SARS-CoV-2 using circular polymerase extension reaction"

(A)

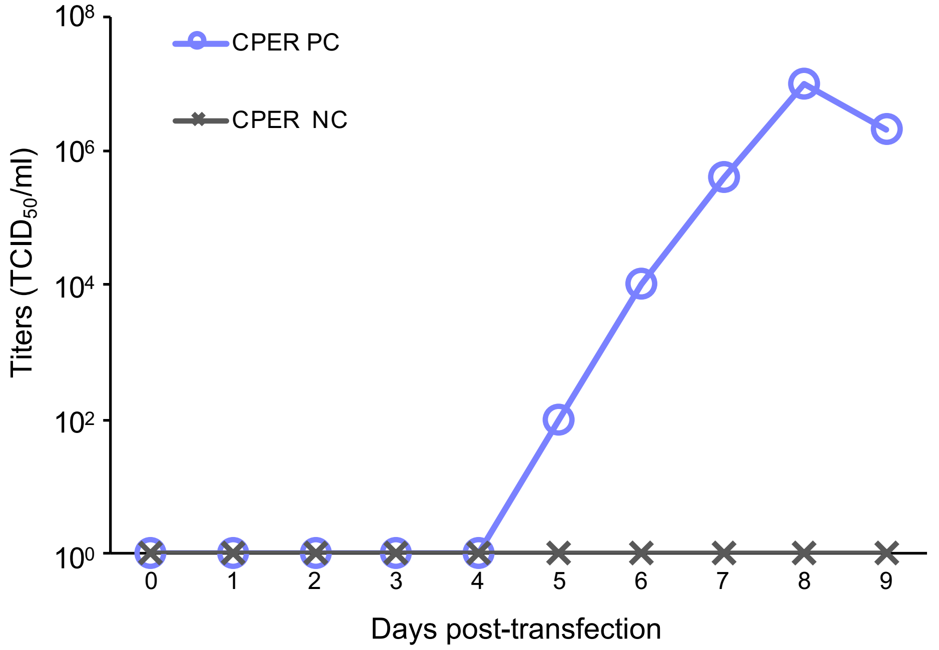


(B)

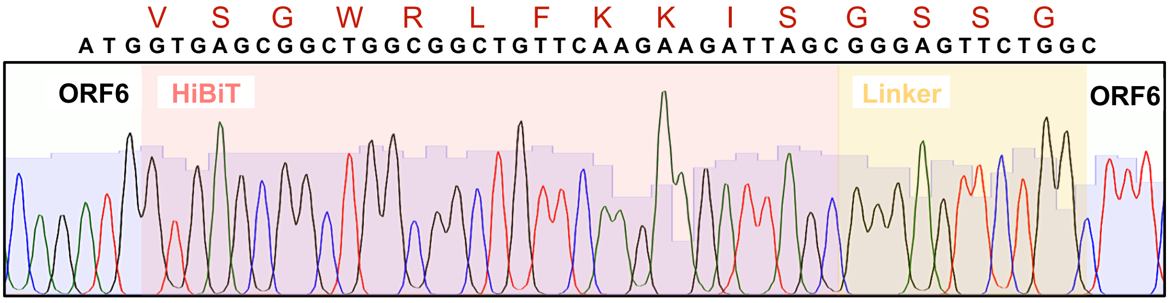


**Figure S1. Characterization of recombinant SARS-CoV-2.**
(A) Time course analysis of infectious SARS-CoV-2 production. CPER products were transfected into HEK293-3P6C33 cells (CPER PC) and infectious titers in the culture supernatants were measured at the indicated time points. As a negative control, the CPER product obtained without fragment F9/10 was transfected into cells (CPER NC). (B) Sequence analysis of the recombinant virus possessing the HiBiT gene in ORF6. The HiBiT gene and a linker sequence were inserted into the N terminus of the ORF6 sequence.

Fig. S1 Torii et al.

Table S1. SARS-CoV-2 DNA fragments used for CPER reaction

| SARS-CoV-2 | Nucleotide no. | bp |
|---|---|---|
| F1 | 1–3,527 | 3,527 |
| F2 | 3,479–6,433 | 2,955 |
| F3 | 6,383–9,811 | 3,035 |
| F4 | 9,360–13,468 | 4,059 |
| F5 | 13,366–16,222 | 2,857 |
| F6 | 16,173–19,300 | 3,128 |
| F7 | 19,247–21,600 | 2,354 |
| F8 | 21,544–25,324 | 3,781 |
| F9/10 | 25,275–29,895 | 4,620 |
| Linker | 1–25 and 29,853–29,895 | 1,175 |

Table S2. Mutations of recombinant SARS-CoV-2 (P0–P2 viruses)

| Sample ID | Passage no. | Nucleotide position | Original | Recombinant | Region |
|---|---|---|---|---|---|
| #1 | P0 | 29,687 | a | t/a | 3'UTR |
| | P1 | 29,687 | a | t/a | 3'UTR |
| | P2 | 29,687 | a | t/a | 3'UTR |
| #2 | P0 | 29,687 | a | t/a | 3'UTR |
| | P1 | 18,490 | c | t | ORF1b |
| | | 23,585–23,599 | deletion | | S1/S2 cleavage site |
| | | 29,687 | a | t/a | 3'UTR |
| | P2 | 6,983 | c | g | ORF1a |
| | | 18,490 | c | t | ORF1b |
| | | 23,585–23,599 | deletion | | S1/S2 cleavage site |
| | | 29,687 | a | t/a | 3'UTR |
